# Exploring chemotaxis in spatio-temporal stoichiometric models of metabolism: *Pseudomonas simiae* as a case study

**DOI:** 10.1101/2025.08.05.668800

**Authors:** Hui Shi, Ercole LiPuma, Haroon Qureshi, Phineas McMillan, Jing Zhang, Melisa Osborne, Ilija Dukovski, Daniel Segrè

**Affiliations:** Biochemistry & Molecular Biology Program, Boston University, Boston, MA, USA; Bioinformatics Program, Faculty of Computing and Data Sciences, Boston University, Boston, MA, USA; Boston University Academy, Boston, MA, USA; Biological Design Center, Boston University, Boston, MA, USA; Center for Advanced Interdisciplinary Research, Ss. Cyril and Methodius University, Skopje, N. Macedonia; Department of Physics, Boston University, Boston, MA, USA; Department of Biomedical Engineering, Boston University, Boston, MA, USA; Department of Biology, Boston University, Boston, MA, USA

**Keywords:** Microbial Communities, Chemotaxis, Flux Balance Analysis, Dynamic Flux Balance Analysis, Spatio-Temporal Simulation, COMETS, Genome Scale Models

## Abstract

Chemotaxis is the movement of cells or organisms in the direction of an increasing or decreasing gradient of a chemical, i.e. a chemoattractant or chemorepellent, respectively. In bacteria, this movement is typically manifested through the presence of bias in the random walk generated by run-and-tumble motility. Often, the chemoattractant is consumed by the organism, thus coupling the biophysical, directed motion with the biochemical, metabolic activity in the growing bacterial colony. In this chapter, we describe our approach to simulating chemotaxis, coupled with detailed metabolic activity, within the framework of our modeling software COMETS (Computation of Microbial Ecosystem in Time and Space). By combining genome-scale metabolic models, dynamic flux balance analysis and the newly implemented biophysical chemotaxis model, we built and qualitatively tested a chemotaxis module for COMETS. We applied this model to an automatically constructed genome-scale model of the plant commensal bacterium *Pseudomonas simiae WCS417*, which we hypothesized would perform chemotaxis on amino acids similar to its close relative *Pseudomonas putida*. COMETS simulations recapitulated the typical ring colony pattern, which is due to propagation at the colony front, indicative of bacteria growing on a chemoattractant-filled plate. By allowing for detailed examination of spatial distributions and time dynamics of the biomass, as well as concentration profiles of the metabolites involved in the growth and chemotaxis in bacterial colonies, the software method described here enables simulations of complex microbial patterns involving metabolism and directed motility. Beyond testing the appearance of the ring, future model validation could include comparisons of local growth rates and metabolic fluxes, including rates of metabolite uptake and secretion. The newly added COMETS chemotaxis capability integrates flux balance modeling with gradient-induced biomass propagation, paving the way for more realistic simulations of microbial ecosystem dynamics in complex environments, such as plant roots, where chemotaxis may play a fundamental role in microbiome-host dynamics.

## Introduction

Beyond displaying dynamic taxonomic compositions and metabolic capabilities, microbiomes can be viewed as complex biophysical systems whose spatial architecture is strongly coupled with their functions. These biophysical and structural properties can affect the role of metabolism in the human microbiome, as well as in marine, soil and plant-associated microbial communities [1], [2], [3], [4]. Much work has been done to study the spatio-temporal dynamics of bacterial colonies and communities, i.e. the way populations of bacteria arrange into macroscopic spatial patterns that can change in time. These studies range from spatial scales as small as Petri dishes [5], [6], [7], [8], [9], [10] to scales as large as complex root microbiomes [11] or even whole lakes [12]. For example, root microbiomes are composed of bacteria that can benefit from plant root exudates at different locations and secrete compounds beneficial to the plant [13]. The knowledge of spatially structured interactions between the bacteria and the plant through exchange and sensing of these compounds could provide meaningful insights into rhizosphere microbiome dynamics, with important consequences for agricultural sustainability and biogeochemical cycles.

Predicting microbial dynamics in such spatially structured systems requires an integrated understanding of the interplay between metabolic exchanges and biophysical processes related to the growth and spread of microbial populations. In prior work, we have developed a software package called COMETS (Computation of Microbial Ecosystem in Time and Space) [14], [15] which simulates the growth of bacterial colonies by combining a dynamic version of flux balance analysis (FBA) with biophysical models of cellular diffusion, motility and demographic fluctuations. FBA, which has been described in detail before [16], [17], [18], [19], [20], generally relies on two major assumptions: (i) that the metabolic network is at steady state under a given environmental condition and (ii) that the network has evolved towards the optimization of a given goal, captured mathematically in a linear objective function. Under these assumptions, FBA is expressed in the form of a convex optimization problem that can be solved using linear programming, yielding predictions of the biomass production rate (or growth rate of the cell), as well as intracellular and exchange (transport) fluxes for all reactions [16], [17], [19], [20], [21]. FBA has been particularly helpful when applied to genome scale models (GEMs), i.e. detailed reconstructions of the metabolic networks of individual organisms, based on enzyme content, encoded in their genomesn[20], [21]. An extension of FBA that is particularly valuable for studying metabolism in microbial communities is dynamic flux balance analysis (dFBA). dFBA still assumes a steady state for intracellular metabolism but treats environmental abundances of different microbes and metabolites as dynamic variables [22], [23], [24], [25]. COMETS can take as input a set of GEMs and environmental conditions and use dFBA and diffusion equations to predict the dynamics of molecules and microbes in structured environments. COMETS is an open-source collaborative platform for modeling microbial ecosystems computationally [14], [15], [26].

Chemotaxis is the oriented (biased) motion of bacteria towards a chemical compound (chemoattractant) or away from it (chemorepellent). It is generally accepted that chemotaxis enables bacteria to sense gradients and move towards resources that help them survive [27], [28], or move away from chemicals that are detrimental [29]. In the absence of chemotaxis or other gradients, bacterial motility is generally observed in the form of an unbiased, random walk (Fig. 1A) [19]. Bacteria perform runs, movement in straight lines, and tumbles, reorientations, often by switching directionality of flagella rotation. In an unbiased random walk, the distribution of run lengths and the frequency of tumbles does not change over time and is not sensitive to the chemical concentrations in the environment. In COMETS, the unbiased movement of bacteria is modeled as diffusion to simulate bacterial movement by swimming [31]. Chemotaxis, too, is a result of a random walk composed of a series of straight runs and random tumbles. However, in the case of chemotaxis, the random walk is biased by the presence of a chemical gradient, as bacteria may sense local differences in the concentration of the chemical and change the frequency of tumbles [31] (Fig. 1B).

**Figure 1:**
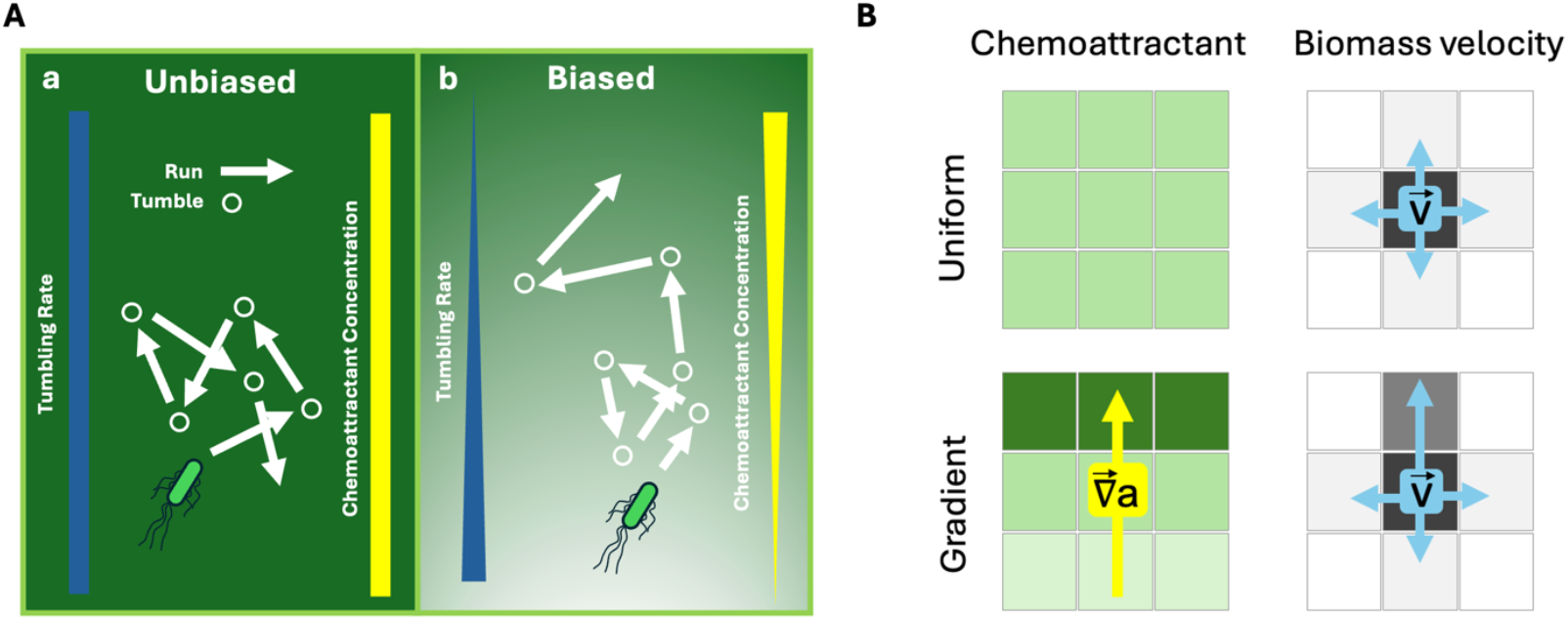
Chemotaxis as biased random walk of bacteria and its implementation in COMETS. **A)** a) Unbiased random walk consistent with normal bacterial motility in media with no gradient. The walk is performed as a stochastic series of straight directed flights (arrows) and sudden changes of direction, tumbles (circles). The average rate (frequency) of the tumbles is constant; b) Biased random walk consistent with chemotaxis, in the presence of a gradient of chemical concentration. When the bacterium senses an increasing concentration of the chemoattractant, the tumble frequency decreases, leading to longer flights, and to overall movement along the chemoattractant gradient. **B)** The biomass propagation in COMETS is modeled by a differential equation with two terms: (i) term for unbiased diffusion and (ii) chemotaxis term proportional to the gradient of the chemoattractant. In the absence of a chemical gradient (top left), the biomass at every step of the numerical algorithm is propagating equally in all directions (top right). In the presence of a chemoattractant gradient (bottom left), the biomass chemotaxis velocity is proportional to the chemical gradient, and its motion is biased in the direction of higher chemoattractant concentration (bottom right).

A variety of mathematical models have been proposed to explain and predict chemotaxis. The Keller-Segel model (KS model) [32], a classical, early model for chemotaxis, was originally developed to describe how slime molds move, and later adapted to model chemotaxis in multiple systems [33],[34],[35],[36]. By explicitly incorporating population growth and nutrient dynamics, a subsequent model, the Growth Expansion model (GE model) [40],[41], enabled more realistic predictions of bacterial chemotaxis, and accurately reproduced the characteristic ring observed at the propagating front of bacterial colonies grown in the presence of a chemoattractant that is metabolized [31],[37],[38],[39]. The GE model is therefore a natural choice for incorporation of a chemotaxis module in COMETS.

The GE model in COMETS is implemented by numerically solving the following equations on a discrete two-dimensional grid (Fig. 1B):

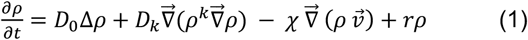

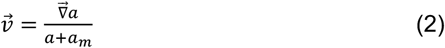

The time dynamics of the local colony biomass, *ρ*, is determined by its diffusion, chemotaxis and local growth (with rate *r*). The first two terms on the right-hand side of Equation 1, 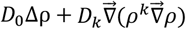, describe the propagation of a bacterial colony, irrespective of chemotaxis, as previously implemented in COMETS [14]. The parameters of this component of the model are the linear diffusivity *D*_0_, the nonlinear diffusivity *D*_*k*_ and the exponent *k*. Here we assume *D*_0_ ≠ 0 and *D*_*k*_ = 0, simulating bacteria that actively swim through simple linear diffusion (i.e., linear with respect to *ρ*). The term 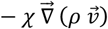 is the one that represents chemotaxis, where *χ* is the chemotaxis coefficient. The velocity of the biomass, calculated in equation (2), is proportional to the gradient of the concentration of the chemoattractant, 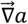. The velocity saturates at a concentration *a*_*m*_, which represents the sensitivity of the chemotactic response. For *a* ≫ *a*_*m*_ the bacteria lose their sensitivity to the gradient and chemotaxis [29, 30]. The last term in equation (1) models the local growth of the biomass.

In this chapter, we will describe in detail how to implement simulations of chemotaxis coupled with genome-scale metabolism and biophysical processes using COMETS. The goal of the specific simulations presented here is to show how COMETS can recapitulate characteristic patterns displayed by bacterial for which a given chemoattractant is also a major nutrient. We will illustrate the process starting from the construction of a genome-scale stoichiometric model for *Pseudomonas simiae WCS417*. The model will be used to implement COMETS simulations of chemotaxis in Petri dishes inoculated in the presence of a chemoattractant, alanine. This specific bacterium was chosen for our case study because it is known to be able to colonize plant roots, promote growth, and provide overall benefits to its host [42]. The output of the simulations is the spatially detailed and time resolved dynamics of the biomass density, concentrations of all extracellular metabolites, as well as all reaction fluxes (rates), including the growth rate and chemoattractant uptake. In addition to describing the results, we will illustrate recommended steps for the visualization and analysis of simulation data.

## Materials

### COMETS and COMETSPy

COMETS (Computation of Microbial Ecosystems in Time and Space) is a software for computational modeling of microbial ecosystem metabolism in structured environments. The current version (2.12.4) is limited to two-dimensional surfaces. COMETS is written in Java and relies on one of two optimizer software packages: Gurobi and Glop. In addition to the core Java code, a Python interface, COMETSPy is available as a user interface (UI) [14]. Detailed information about COMETS, including installation, is available at www.runcomets.org.

Download and install COMETS from https://www.runcomets.org/installation. The source code of COMETS is freely available at https://github.com/segrelab/comets. The COMETSPy source code is available at https://github.com/segrelab/cometspy. The COMETSPy scripts and the genome-scale model used in this chapter can be found at: https://github.com/segrelab/COMETS_Chemotaxis/

### Kbase

Kbase (Knowledgebase) is a software developed by the US Department of Energy [43]. It integrates several methods for systems biology research and data analysis. It can be accessed at: https://www.kbase.us/. Datasets and apps used for this work are available in the form of the following Kbase Narrative: https://narrative.kbase.us/narrative/222319

### Software requirements and the procedure for installing COMETS

COMETS is written in Java. Obtain and install Java from https://www.java.com/en/download/manual.jsp

### Python

The user interface to COMETS, COMETSPy, is written in Python. Although COMETS can be used without it, we recommend using COMETSPy with Jupyter notebooks, which have a convenient user interface. We recommend installing and using Python from the Anaconda distribution available at: https://www.anaconda.com/download

### CobraPy

CobraPy is a Python package that provides an interface to constraint-based metabolic modeling. We used this package to facilitate the handling of our genome-scale models. CobraPy can be downloaded and installed using the pip command in an Anaconda power shell terminal or a command line terminal: `pip install cobra` or alternatively from: https://opencobra.github.io/cobrapy/

### Pandas

Pandas is a Python package that enables users to manipulate data in the form of data frames. We use this package to handle and analyze output data (biomass, concentration, etc.) from simulation runs. Pandas can be downloaded and installed using the pip command: `pip install pandas` or alternatively from: https://pandas.pydata.org/docs/getting_started/install.html

### Linear optimization software

The Flux Balance Analysis component of COMETS requires running several steps of linear programming (LP) optimization. In COMETS, the following two optimizers can be used:

### Glop

The default linear optimizer in COMETS is Glop. It is part of the OR-Tool suite: https://developers.google.com/optimization. Glop is distributed with COMETS and the user does not need to perform any additional installation steps.

### Gurobi

An alternative optimizer that can be used in COMETS is Gurobi. The package can be downloaded and installed from http://www.gurobi.com/. The installation of Gurobi requires obtaining a license from http://www.gurobi.com/downloads/licenses/license-center. Academic users that will use COMETS on an individual basis may obtain a free academic license. When the installation of Gurobi is finished, it is very important to have the environment variable GUROBI_HOME set to the directory where Gurobi was installed. In Windows, this variable is set automatically during the installation process. In Linux and MacOS, however, depending on the system, sometimes this is not the case. It is therefore important to make sure that a line such as the following example: export GUROBI_HOME=/usr/gurobi/gurobi902/linux64/is included in the user’s.bashrc file in Linux, or the corresponding file for an alternative shell. The version name and number in gurobi902 should be set to the one installed.

Here, we use Gurobi version 9.0.2 (i.e. ‘gurobi902’) as a representative example version of Gurobi, but we anticipate that the following steps will continue to be valid for subsequent implementations.

In MacOS, the line:

`export GUROBI_HOME=/Library/gurobi902/mac64/`

should be included in the.bash_profile file in older versions of MacOS, or.zshrc in the latest version of MacOS. The version name and number in gurobi902 should be set to the one installed.

## Methods

### Complete pipeline from genome to chemotaxis simulation in COMETS

1. Construction of a draft genome-scale metabolic model of *Pseudomonas simiae* WCS417 in Kbase (https://narrative.kbase.us/narrative/222319)

1.1 Download the genome of the strain *P. simiae* WCS417 from NCBI: https://www.ncbi.nlm.nih.gov/nuccore/NZ_CP007637.1

Click on “Assembly” link in the bar on the right-hand side. This will take you to: https://www.ncbi.nlm.nih.gov/datasets/genome/GCF_000698265.1/

Click on “Download.” A window will pop up to choose the sources and types of data to download. On the popup window, choose “All” sources and “Genome sequences (FASTA),” “Sequence and annotation (GBFF)” and “Protein (FASTA).” Click on “Download.” Unzip the downloaded files.

1.2 Create an account and login to Kbase: https://narrative.kbase.us

Click on “Navigator” in the top left corner. Create a new narrative by clicking on “+ New Narrative” in the top right corner. Once a new narrative is created, rename it to your liking by clicking on “Unnamed” in the top left corner. In this case, name it “P. simiae WCS417.”

1.3 Click on “Add Data” on the “DATA” panel (stage) on the left (see Fig. 2). On the popup panel, click “Import” in the top right. Click on “Select” and select the file “genomic.gbff” from the genome folder that you downloaded in step 1.1.

**Figure 2:**
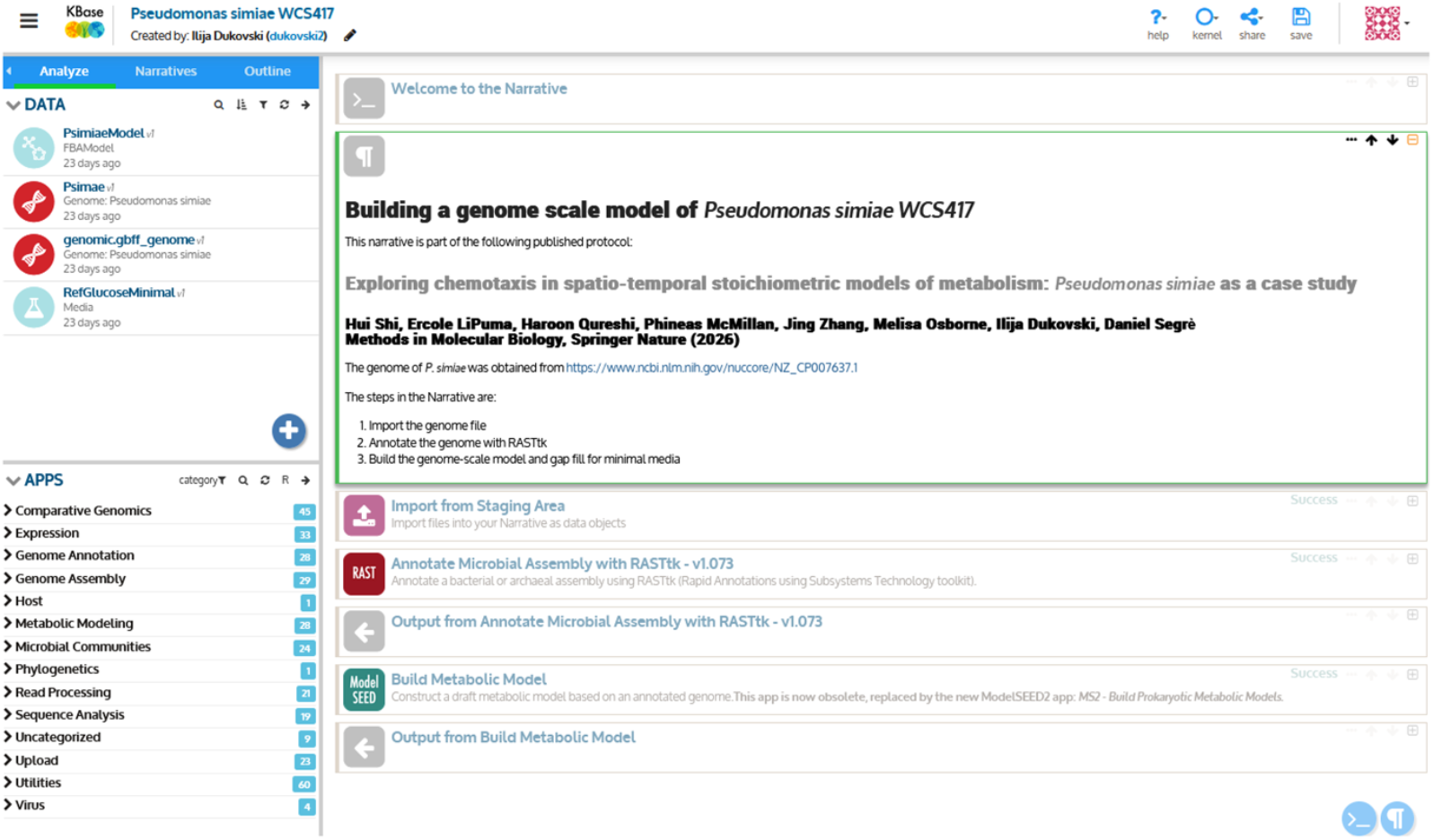
Kbase Narrative for building a genome-scale metabolic stoichiometric model of *P. simiae*. The Kbase Narrative for building genome-scale metabolic model of *P. simiae* consists of three applications: Import from Staging Area, Annotate Microbial Assembly with RASTtk and Build Metabolic Model. The imported data as well as the model output are in the Data panel on top left. The apps are loaded from the Applications panel on the bottom left.

The full path is: ncbi_dataset\ncbi_dataset\data\GCA_000698265.1\genomic.gbff.

The app “Import from Staging Area” will be added to your Narrative. Choose “GenBank Genome” in the “Import as” field and type “P. simiae” in “Name” and click on “Run.” This will import the file, and the genome and assembly files will show up on the left “DATA” area.

1.4 Add minimal media to your data, by clicking on the plus sign in the left panel. Click on “Public” on the top and choose “Media” as data type and search for “RefGlucoseMinimal.” Choose “RefGlucoseMinimal” and click on “Add” arrow. This will add “RefGlucoseMinimal” media to your data.

1.5 Open the “Annotate Microbial Assembly with RASTtk” app from the “Genome Annotation” app block. This will add the app to your Narrative.

1.6 Choose the file “genomic.gbff_genome” file. Add the scientific name P. simiae. Add the name of the output file. In this case, it is “Psimiae.” Press “Run.” This will create a genome annotation file.

1.7 Open the “Metabolic Modeling” block in the Apps area on the bottom left, and choose the “Build Metabolic Model” app. This will add the app to your Narrative.

1.8 In the “Build Metabolic Model” app, choose your genome annotation and media. Use “Automatic selection” for “Template reconstruction.” Choose “Aerobic glucose” for “Media for core ATP gapfilling”. Select “Gapfill model?” Choose a name for your model. In this case, it is “PsimiaeModel.” Click on “Run.” This will build your model.

1.9 Download the model file from Kbase. On the “DATA” panel on the left, click on the model “PsimiaeModel.” Choose the “export/download data” button and download the model in SBML format. Rename the model as needed, in this case Psimiae_withATPM.xml.

2. COMETS Simulations of a *P. simiae* colony propagation with and without chemotaxis to alanine (Psimiae_chemotaxis_alanine.ipynb)

2.1. Download the scripts and all relevant files from https://github.com/segrelab/COMETS_Chemotaxis.

2.2 Start Jupyter Notebook, navigate to COMETS_Chemotaxis/simulation_notebooks/and open the Jupyter notebook Psimiae_chemotaxis_alanine.ipynb.

2.3 Import the relevant python libraries. These libraries can be downloaded using “pip install” and the package name in terminal. They can also be installed with pip as shown in the first line. See the “Materials” section for more information.

~~~
#!pip install cobra
# Install cobra if have not done it yet.
import cometspy as c
from matplotlib import pyplot as plt
import pandas as pd import numpy as np
#Import cobra libraries from the cobra toolbox.
#They are needed to load model files in sbml or other formats.
from cobra.io import load_json_model, save_json_model, load_matlab_model, save_matlab_model, read_sbml_model, write_sbml_model
~~~

2.4 Load the genome scale model (GEM) of the organism we want to model.

The argument should be the name of the file, or full path if not in the working directory. See Notes for more information about loading GEMs.

~~~
# First, load the genome-scale model of P. simiae
cobra_model = read_sbml_model(“Psimiae_withATPM.xml”)
~~~

2.5 Set the maintenance reaction lower bound. The maintenance reaction lower bound, i.e. ATP maintenance requirement needs to be specified as needed. Set it to zero or as needed.

~~~
#Modify the ATP maintenance lower limit as needed.
#In this example we will not impose any ATP maintenance limit.
cobra_model.reactions.get_by_id(“ATPM”).lower_bound = 0.0
~~~

2.6 Create a COMETS format metabolic model from the SBML Cobra model format. Use the command open exchanges to ensure that all exchange lower bounds are set to −1000.

~~~
#Create a COMETS model from the SBML/Cobra model format.
Psimiae = c.model(cobra_model)
Psimiae.id = “psimiae”
Psimiae.open_exchanges() # Make sure that exchanges are active.
~~~

2.7 Specify the parameters of the different terms in the biophysical model of colony biomass propagation (Eq. 1). In particular, it is important to set parameters for the first two terms in Eq, 1, representing standard nonbiased diffusion of bacteria, and the third term, describing chemotaxis. See Notes for detailed description of the input parameters.

~~~
#Set the parameters of the biomass propagation model.
#In this case, we set the parameter for simple, linear diffusion of the biomass.
#This setting corresponds to active swimming bacteria. Psimiae.add_nonlinear_diffusion_parameters(D0 = 1.0e-7, Dk = 0.0, exponent = 0.0, hilln = 1.0, hillk = 0.0)
#Set the parameters of the chemotaxis model. Psimiae.add_nonlinear_diffusion_chemotaxis_parameters(hilln = 1.0, hillk = 0.0)
~~~

2.8 Define the array that holds information about the initial amount and location of the bacteria. Users should consider a set of coordinates to serve as the midpoint of the circle and then determine the size of the circle. The current code defines (100, 100) to be the midpoint with a circle of radius 5 grid points. Each grid point within the circle is initialized with 1e-7 grams of biomass.

~~~
*# Create an initial inoculum with P. simiae, that will be added to the
spatial layout in the next cell*.
Psimiae.initial_pop **=** []
**for** i **in** range(95,105):
     **for** j **in** range(95,105):
                Psimiae.initial_pop.append([i,j,1e-7])
~~~

2.9 Set the desired size of the grid and layout.

~~~
*# Create a spatial layout, a square of 201 by 201 grid points, with the P. simiae inoculum created above*.
*# Later we will define each grid point to have a linear length of 0*.*03 cm*.
*# We are thus creating a 6 by 6 cm, two-dimensional Petri dish layout*.
layout **=** c.layout([Psimiae])
layout.grid **=** [201, 201]
~~~

2.10 Define the amount of each compound to be used in the medium. See Notes for more information about compound naming conventions.

~~~
*# Pour the media in the layout*.
*# The model of P. simiae created in Kbase uses ModelSeed names for the metabolites*.
*# If you want to add additional metabolite, you must find its ModelSeed name*.
*# Here the unit is mmols per grid pixel in the layout*.
layout.set_specific_metabolite(‘cpd00013_e0’, 0.0000029)*#NH3*
layout.set_specific_metabolite(‘cpd00099_e0’, 1000) *#Cl*
layout.set_specific_metabolite(‘cpd00205_e0’, 1000) *#K*
layout.set_specific_metabolite(‘cpd00971_e0’, 1000) *#Na*
layout.set_specific_metabolite(‘cpd00067_e0’, 1000) *#H*
layout.set_specific_metabolite(‘cpd00009_e0’, 0.0000027) *#H2PO4*
layout.set_specific_metabolite(‘cpd00254_e0’, 1000) *#Mg*
layout.set_specific_metabolite(‘cpd00048_e0’, 0.000018) *#SO4*
layout.set_specific_metabolite(‘cpd00030_e0’, 1000)*#Mn2+*
layout.set_specific_metabolite(‘cpd00001_e0’, 1000) *#H2O*
layout.set_specific_metabolite(‘cpd10515_e0’, 1000)*#Fe+2*
layout.set_specific_metabolite(‘cpd00063_e0’, 1000) *#Ca*
layout.set_specific_metabolite(‘cpd00149_e0’, 1000)*#Co2+*
layout.set_specific_metabolite(‘cpd00034_e0’, 1000) *#Zn*
layout.set_specific_metabolite(‘cpd00058_e0’, 1000) *#Cu*
layout.set_specific_metabolite(‘cpd09225_e0’, 0.00000010) *#H3BO3*
layout.set_specific_metabolite(‘cpd11574_e0’, 0.000000076) *#MoO4*
layout.set_specific_metabolite(‘cpd00244_e0’, 1000)*#Ni2+*
layout.set_specific_metabolite(‘cpd15574_e0’, 0.000000048) *#WO4*
layout.set_specific_metabolite(‘cpd00104_e0’, 1000) *#biotin*
layout.set_specific_metabolite(‘cpd00393_e0’, 1000) *#folic acid*
layout.set_specific_metabolite(‘cpd00263_e0’, 1000) *#pyridoxine*
layout.set_specific_metabolite(‘cpd00220_e0’, 1000) *#riboflavin*
layout.set_specific_metabolite(‘cpd00305_e0’, 1000) *#thiamine*
layout.set_specific_metabolite(‘cpd00218_e0’, 1000) *#nicotinic acid*
layout.set_specific_metabolite(‘cpd00644_e0’, 1000) *#pantothenic acid*
layout.set_specific_metabolite(‘cpd03424_e0’, 1000) *#vitamin b12*
layout.set_specific_metabolite(‘cpd00443_e0’, 1000) *#4-Aminobenzoate*
layout.set_specific_metabolite(‘cpd00541_e0’, 1000) *#thioctic acid (lipoate)*
layout.set_specific_metabolite(‘cpd00027_e0’, 2e-5) *#D-glucose #1e-5*
layout.set_specific_metabolite(‘cpd00007_e0’, 1000) *#O2*
layout.set_specific_metabolite(‘cpd00035_e0’, 1e-5) *#alanine*
~~~

2.11 Assign the chemotaxis coefficient and uptake parameters to the chemoattractant. See notes for the description of the input parameters.

~~~
*# Set the chemotaxis model parameters. Here we define alanine (cpd00035_e0 is #alanine ModelSeed name) as chemoattractant*.
layout.set_chemotaxis(0, “cpd00035_e0”, 0.0001, 1e-6)
~~~

2.12 Define the diffusivity of the media.

~~~
*# Set the diffusivity of the media. The unit is cm^2/sec*.
layout.set_metabolite_diffusion(5e-6)
~~~

2.13 Modify the parameters of the simulation as needed. See Notes for a detailed description of the parameters.

~~~
*#Now create a class of parameters for the simulation*.
parameters **=** c.params()
*#Populate the parameters with values*.
parameters.all_params[“maxCycles”] **=** 500 *#The total numer of cycles to be
run*.
parameters.all_params[“timeStep”] **=** 0.05 *# Unit hours*.
parameters.all_params[“spaceWidth”] **=** 0.03 *# Unit cm. Linear size per layout grid pixel*.
parameters.all_params[“biomassMotionStyle”] **=** “Nonlin Diff Chemotaxis 2D”
*#The name of the biophysical model*.
parameters.all_params[“defaultKm”] **=** 0.069 *# Michaelis constant for the uptake. Unit mmol/cm^3*.
parameters.all_params[“defaultVmax”] **=** 7. 1 *# Maximum uptake flux. Unit mmol/gh*.
parameters.all_params[“maxSpaceBiomass”] **=** 10 *# This is a large number to stop growth if the biomass surpasses it. Unit grams*.
parameters.all_params[“minSpaceBiomass”] **=** 1.e-11 *# If the biomass falls below this value, it will be reset to zero*.
parameters.all_params[“numRunThreads”] **=** 4 *# Numer of parralel threads to run in the simulation*.
parameters.all_params[“writeBiomassLog”] **= True** *# Save the biomass spatio-temporal data*.
parameters.all_params[“BiomassLogRate”] **=** 50 *# The cycles period to save the biomass data. This will save avary 50th time step*.
parameters.all_params[“writeTotalBiomassLog”] **= True** *# Save the total integrated biomass data*.
parameters.all_params[“totalBiomassLogRate”] **=** 1 *# The cycles period to save the total biomass data*.
parameters.all_params[“writeFluxLog”] **= True** *# Save all the reaction fluxes spatio-temporal data*.
parameters.all_params[“FluxLogRate”] **=** 50 *# The cycles period to save the fluxes data*.
parameters.all_params[“writeMediaLog”] **= True** *# Save all the reaction media (extracellular metabolites) spatio-temporal data*.
parameters.all_params[“MediaLogRate”] **=** 50 *# The cycles period to save the media data*.
~~~

2.14 Consolidate the layout and parameters into a COMETS object and then run the simulation.

~~~
*#Put together the layout and the simulation parameters in a single comets #simulation*.
simulation **=** c.comets(layout, parameters)
*#Run the simulation. The argument False provides that the output files are not #deleted after the simulation*.
*#Setting the parameter to True, will clean the working directory clean of all #output*.
simulation.run(False)
~~~

2.15 Print the output of the simulation for progress monitoring and debugging purposes if needed.

~~~
*#Check the command line output to confirm that the run finished with no
#errors*.
print(simulation.run_output)
~~~

## 3. Visualization and analysis of the simulation output

### Biomass

### 3.1 Define the function that will help with biomass visualization

~~~
*#This function will load the file and return a matrix of spatial biomass
#distribution at a given time*
import numpy as np
def read_biomass_file_and_process(filename, Nx, Ny, time_cycle):
    “““
    Reads a file with the specified format and processes each line.
    Args:
       filename: The path to the file to be read.
       Nx: Size of the spatial grid in x direction.
       Ny: Size of the spatial grid in y direction.
       time_cycle: The simulation time step to be loaded.
   “““
matrix_data=np.zeros((Nx+1,Ny+1))
*#Create an empty matrix with the spatial grid size for the biomass data*.
*# Read the file and select the biomass grid content at the time_cycle*.
with open(filename, ‘r’) as file:
   for line in file:
    line = line.strip()
    if not line:
       continue
    values = line.split()
    try:
       val1 = int(values[0]) *#time cycle*
       val2 = int(values[1]) *#x coordinate*
       val3 = int(values[2]) *#y coordinate*
       string_val = values[3] *#name of the model*
       scientific_val = float(values[4]) *#biomass content in grams
       #select the specified time cycle and populate the matrix with biomass
content*
       if val1==time_cycle:
              matrix_data[val2,val3]=scientific_val
       except (ValueError, IndexError) as e:
           print(f”Error processing line: {line} - {e}”)
       return matrix_data
~~~

### 3.2 Load in the biomass file. See notes for details on input file name

~~~
*# Now we will load the biomass file*.
*# Change the filename string as it was produced by the simulation*
filename **=** “biomasslog_0×14c6a7706410”
biomass_data**=**read_biomass_file_and_process(filename, Nx**=**201, Ny**=**201, time_cycle**=**250)
~~~

### 3.3 Convert the units in the original biomass file to our preferred version

~~~
*#In this step we will transform the data from biomass content per pixel in
#grams*,
*#to biomass density per cm^2. The linear size of a pixel in this simulation
#was L=0*.*03cm*.
*#The linear size should be changed according to the one set in the
simulation*.
L**=**0.03
biomass_data**=**biomass_data**/**(L**\***L)
~~~

### 3.4 Import relevant libraries and graph the biomass. The results are shown in Figs. 3B and 3E

~~~
*#Now we are ready to visualize the biomass*.
import matplotlib.pyplot as plt
import matplotlib.patches as patches
fig, ax = plt.subplots()
*# Display the data as an image and get the mappable object*.
*# The cmap (colormap) was chosen to produce a grayscale image*.
im = ax.imshow(biomass_data, cmap=‘Greys_r’)
*# Create the colorbar*
cbar = fig.colorbar(im)
*# Set the title of the colorbar*
cbar.ax.set_title(‘[g/cm\u00b2]’, loc=‘center’) *# loc=‘center’ is default but
can be specified*
plt.axis(‘off’) *#do not plot the axis
# Show a size bar with a “1cm” label*
plt.text(15, 180, ‘1 cm’, color=‘white’, fontsize=12)
*# Define rectangle parameters: (x_lower_left, y_lower_left, width, height)*
rect_x = 10
rect_y = 185
rect_width = 33
rect_height = 2
*# Create the Rectangle patch*
rect = patches.Rectangle((rect_x, rect_y), rect_width, rect_height,
linewidth=2, edgecolor=‘w’, facecolor=‘w’)
*# Add the patch to the Axes*
ax.add_patch(rect)
*#Add title*
plt.title(“Biomass density”)
*# First save the image, then display the plot*
plt.savefig(“biomass.png”) plt.show()
~~~

**Figure 3:**
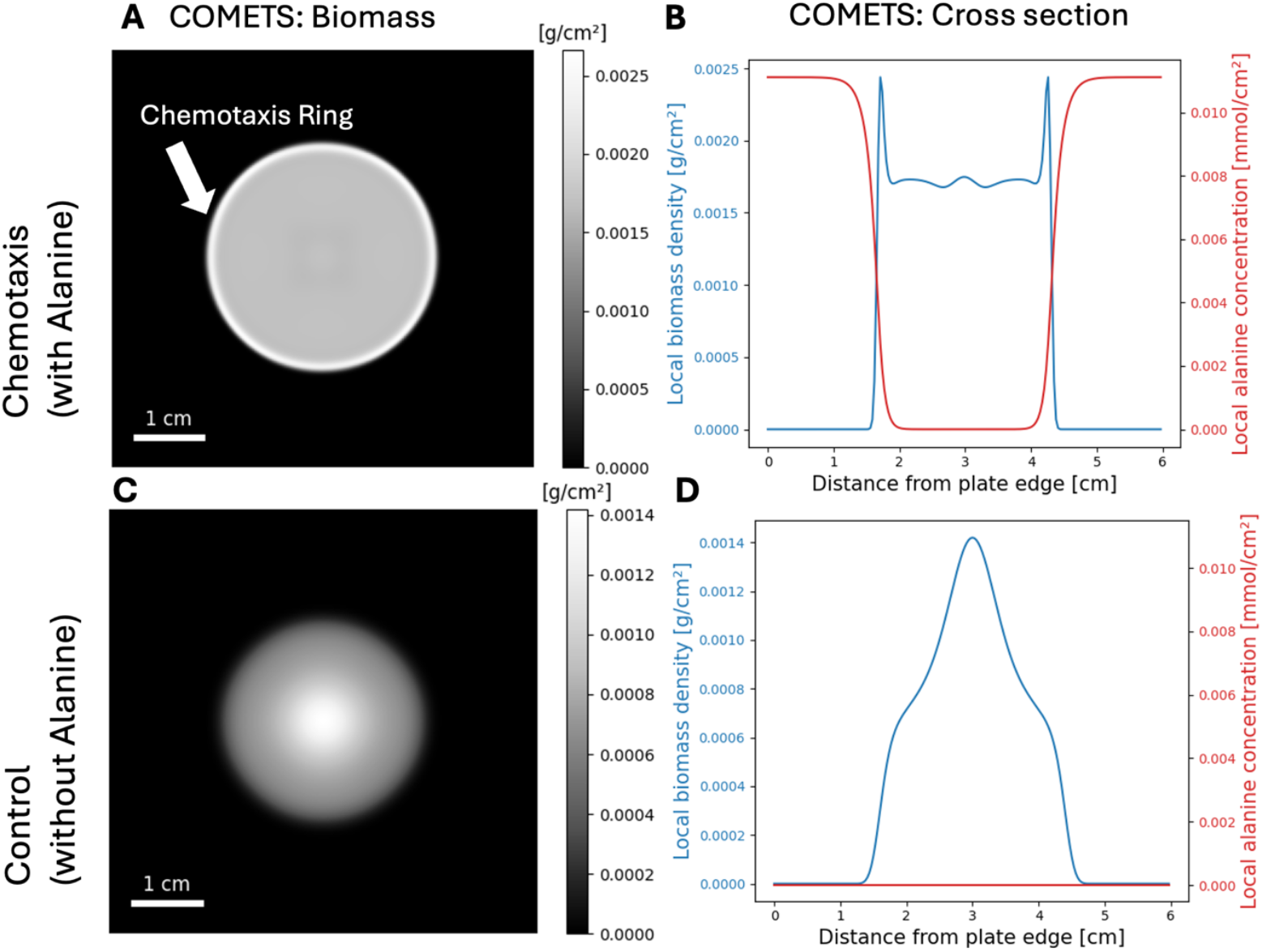
COMETS simulations of chemotaxis of *P. simae* in the presence of alanine, compared to control (no alanine). **A)** Simulated *P. simiae* colonies, grown in the presence of alanine [40], [41]. The previously documented chemotaxis ring at the edge of the colony [40], [41] is evident in the simulated colonies. **B)** The cross section of the simulated colony, with the leading peak coinciding with the gradient of alanine, created due to the consumption of alanine by the bacteria. **C)** A simulation experiment without alanine in the growth medium. The chemotaxis ring is absent in this case. **D**) The cross section of the simulated colony without alanine. No leading peak in the cross section can be seen in this condition.

### Media

### 3.5 Define the function that will help with media visualization

~~~
*#This function will load the file and return a matrix of spatial biomass #distribution at a given time*
def read_media_file_and_process(filename,metabolite, Nx, Ny, time_cycle):
   “““
   Reads a file with the specified format and processes each line.
   Args:
      filename: The path to the file to be read.
      metabolite: The name of the metabolite data to be loaded.
      Nx: Size of the spatial grid in x direction.
      Ny: Size of the spatial grid in y direction.
      time_cycle: The simulation time step to be loaded.
  “““
matrix_data=np.zeros((Nx+1,Ny+1)) *#Create an empty matrix with the spatial
#grid size for the media data*.
*# Read the file and select the media grid content at the time_cycle*.
with open(filename, ‘r’) as file:
    for line in file:
      line = line.strip()
      if not line:
         continue
      values = line.split()
      try:
         string_val = values[0] *# Name of the metabolite*
         val1 = int(values[1]) *# time cycle*
         val2 = int(values[2]) *# x coordinate*
         val3 = int(values[3]) *# y coordinate*
         scientific_val = float(values[4]) *# metabolite content (in mmol)*
         if string_val==metabolite and val1==time_cycle:
         matrix_data[val2,val3]=scientific_val
         except (ValueError, IndexError) as e:
            print(f”Error processing line: {line} - {e}”)
return matrix_data
~~~

### 3.6 Filter the media file for the media of interest and load the data. See notes on how to handle large files

~~~
*# Now load the data. In this case, the file was too large, so we filtered it
#only for alanine
# with the following command in a terminal:
# cat medialog_0×14c6a7706410* |*grep cpd00035_e0 >alanine
# The input file is named ‘alanine’ in this case*
filename **=** “alanine”
media_data**=**read_media_file_and_process(filename,metabolite**=**‘cpd00035_e0’,Nx**=**2
01,Ny**=**201,time_cycle**=**250)
~~~

### 3.7 Calculate the concentration of alanine based on the amount listed in the files

~~~
*# As for the biomass, we will calculate the concentration of alanine*
L**=**0.03 *#cm*
media_data**=**media_data**/**(L**\***L)
~~~

### 3.8 Import pertinent libraries and visualize the media. The result is shown in Fig. 4A

~~~
import matplotlib.pyplot as plt
import matplotlib.patches as patches
import seaborn
fig, ax **=** plt.subplots()
*# Display the data as an image and get the mappable object*
im **=** ax.imshow(media_data[1:201,1:201], cmap**=**‘viridis’)
*# Create the colorbar*
cbar **=** fig.colorbar(im)
*# Set the title of the colorbar*
cbar.ax.set_title(‘[mmol/cm\u00b2]’, loc**=**‘center’) *# loc=‘center’ is default
but can be specified
#Do not display the axis*
plt.axis(‘off’)
*# Include a size bar and a label*
plt.text(15, 180, ‘1 cm’, color**=**‘black’, fontsize**=**12)
*# Define rectangle parameters: (x_lower_left, y_lower_left, width, height)*
rect_x **=** 10
rect_y **=** 185
rect_width **=** 33
rect_height **=** 2
*# Create the Rectangle patch*
rect **=** patches.Rectangle((rect_x, rect_y), rect_width, rect_height,
linewidth**=**2, edgecolor**=**‘k’, facecolor**=**‘k’)
*# Add the patch to the Axes*
ax.add_patch(rect)
*#Add title*
plt.title(“Alanine concentration”)
*# Save the image*
plt.savefig(‘alanine.png’)
*# Display the plot*
plt.show()
~~~

**Figure 4:**
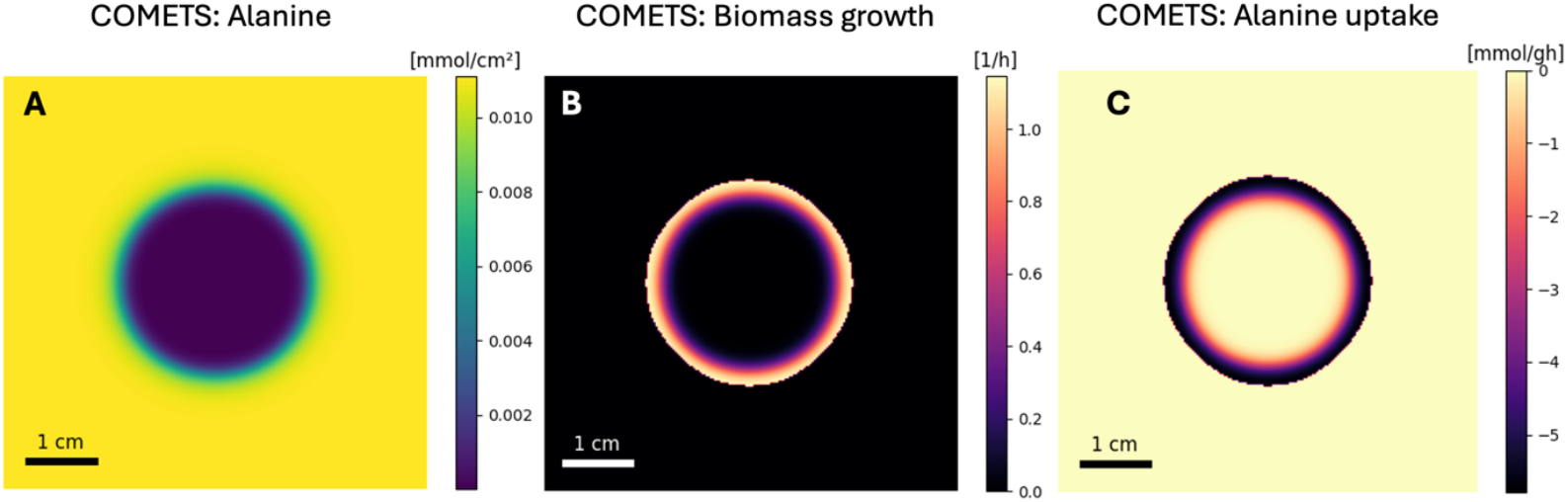
A typical COMETS output consists of a time series of all spatial distributions of metabolite concentrations and reaction fluxes, including biomass growth rate. **A)** Simulated alanine concentration showing the depletion region at the center of the layout and its gradient at the colony edge. **B)** The biomass growth is localized at the edge of the colony and coincides with **C)** the region of alanine uptake. The growth region where the uptake is taking place is where the alanine gradient is produced, which in turn triggers chemotaxis and produces the leading biomass ring, shown in Fig. 3.

### Cross Section of Biomass and Alanine

### 3.9 Define the x coordinate and the range of y coordinates from which to take a cross section

~~~
*# Here we take a cross section in the middle of the layout*
media_cross_section**=**media_data[100,1:201]
biomass_cross_section**=**biomass_data[100,1:201]
~~~

### 3.10 Plot the cross sections. The results are shown in Figs. 3C and 3F

~~~
*# Now we can plot the cross section profile*
import matplotlib.pyplot as plt
import numpy as np
*# Get the Data first*
x**=**np.arange(0.0,6.0,0.03) *#This is the x distance from the plate edge*
y1**=**biomass_cross_section[0:200] *#Biomass cross section*
y2**=**media_cross_section[0:200] *#Alanine cross section
# Create the first plot (left y-axis)*
fig, ax1 **=** plt.subplots()
color **=** ‘tab:blue’
ax1.set_xlabel(‘Distance from plate edge [cm]’, fontsize**=**15)
ax1.set_ylabel(‘Local biomass density [g/cm\u00b2]’, color**=**color,
fontsize**=**15)
ax1.plot(x, y1, color**=**color)
ax1.tick_params(axis**=**‘y’, labelcolor**=**color)
*# Create the second plot (right y-axis)*
ax2 **=** ax1.twinx() *# Instantiate a second Axes that shares the same x-axis*
color **=** ‘tab:red’
ax2.set_ylabel(‘Local alanine concentration [mmol/cm\u00b2]’, color**=**color,
fontsize**=**15)
ax2.plot(x, y2, color**=**color)
ax2.tick_params(axis**=**‘y’, labelcolor**=**color)
*# Adjust layout*
fig.tight_layout() *# Adjust layout to prevent labels from overlapping
#Add title*
plt.title(“Biomass and alanine cross section “)
*#Save the plot and display it*
plt.savefig(“biomass_alanine.png”)
plt.show()
~~~

### Uptake Fluxes and Biomass Growth Rate

### 3.11 Define the function that will read the fluxes from the file

~~~
*#Function to read the fluxes output file*
def read_flux_file_and_process(filename,flux,Nx,Ny,model_number,time_cycle):
   “““
   Reads a file with the specified format and processes each line.
   Args:
      filename: The path to the file to be read.
      flux: The name of the reaction flux to be processed.
      Nx: Size of the spatial grid in x direction.
      Ny: Size of the spatial grid in y direction.
      time_cycle: The simulation time step to be loaded.
   “““
   atrix_data=np.zeros((Nx+1,Nx+1))
   with open(filename, ‘r’) as file:
     for line in file:
       line = line.strip()
       if not line:
          continue
       values = line.split()

       try:
       val0 = int(values[0])
       val1 = int(values[1])
       val2 = int(values[2])
       val3 = int(values[3])
       scientific_val = float(values[flux+3])
       if val0==time_cycle and val3==model_number:
          matrix_data[val1,val2]=scientific_val
except (ValueError, IndexError) as e:
   print(f”Error processing line: {line} - {e}”)
 return matrix_data
~~~

### 3.12 Load in the flux data and get an array of alanine uptake flux

~~~
*#Before loading the fluxes data, you need to know what the location of the
#flux of interest in the array of fluxes is*.
*#Here we will load the alanine uptake flux*.
filename **=** “fluxlog_0×149264947d30”
flux_data**=**read_flux_file_and_process(filename,flux**=**1439,Nx**=**201,Ny**=**201,model_n umber**=**1,time_cycle**=**250)
~~~

### 3.13 Load in relevant libraries and visualize the flux. The result is shown in Fig. 4C

~~~
import matplotlib.pyplot as plt
import matplotlib.patches as patches
import seaborn
fig, ax = plt.subplots()
*# Display the data as an image and get the mappable object*
im = ax.imshow(flux_data[1:201,1:201], cmap=‘magma’)
*# Create the colorbar*
cbar = fig.colorbar(im)
*# Set the title of the colorbar*
cbar.ax.set_title(‘[mmol/gh]’, loc=‘center’) *# loc=‘center’ is default but can be specified*
plt.axis(‘off’)
plt.text(15, 180, ‘1 cm’, color=‘black’, fontsize=12)
*# Define rectangle parameters: (x_lower_left, y_lower_left, width, height)*
rect_x = 10
rect_y = 185
rect_width = 33
rect_height = 2
*# Create the Rectangle patch*
rect = patches.Rectangle((rect_x, rect_y), rect_width, rect_height,
linewidth=2, edgecolor=‘k’, facecolor=‘k’)
*# Add the patch to the Axes*
ax.add_patch(rect)
*#Add title*
plt.title(“Alanine uptake”)
plt.savefig(‘Flux_EX_alanine.png’)
*# Display the plot*
plt.show()
~~~

### 3.14 Read the biomass growth rate

~~~
*#Now we can do the same for biomass growth rate*
biomass_growt_rate**=**read_flux_file_and_process(filename,flux**=**1417,Nx**=**201,Ny**=**20 1,model_number**=**1,time_cycle**=**250)
~~~

### 3.15 Load in the relevant libraries and visualize the biomass flux. The result is shown in Fig. 4B

~~~
import matplotlib.pyplot as plt
import matplotlib.patches as patches
import seaborn
fig, ax **=** plt.subplots()
*# Display the data as an image and get the mappable object*
im **=** ax.imshow(biomass_growt_rate[1:201,1:201], cmap**=**‘magma’)
*# Create the colorbar*
cbar **=** fig.colorbar(im)
*# Set the title of the colorbar*
cbar.ax.set_title(‘[mmol/gh]’, loc**=**‘center’) *# loc=‘center’ is default but can be specified*
plt.axis(‘off’)
plt.text(15, 180, ‘1 cm’, color**=**‘black’, fontsize**=**12)
*# Define rectangle parameters: (x_lower_left, y_lower_left, width, height)*
rect_x **=** 10
rect_y **=** 185
rect_width **=** 33
rect_height **=** 2
*# Create the Rectangle patch*
rect **=** patches.Rectangle((rect_x, rect_y), rect_width, rect_height,
linewidth**=**2, edgecolor**=**‘k’, facecolor**=**‘k’)
*# Add the patch to the Axes*
ax.add_patch(rect)
*#Add title*
plt.title(“Biomass growth rate”)
plt.savefig(‘Flux_EX_alanine.png’)
*# Display the plot*
plt.show()
~~~

## Notes

### Notes for building a genome-scale metabolic model of *Pseudomonas simiae* WCS417 in Kbase

1. The quality of a Genome Scale Models (GEMs), i.e. its capacity to accurately represent the metabolic functions of an organism, is strongly dependent on the quality of the sequenced genome and of the annotation of individual gene functions. In bacterial genomes, and especially in non-model organisms, there is often a significant number of genes with unknown functions. This lack of knowledge is reflected in gaps in the initial reconstruction of the metabolic network. Many model-building algorithms, including the one used here, improve the initial reconstruction through gap filling steps, which aim at completing pathways by adding reactions form a database, based on defined criteria and objectives. An important caveat is that without further manual curation, automatically gap-filled algorithms are still likely to contain errors and inaccuracies. In addition to using thoroughly completed and well annotated genomes, readers should be aware of the current limitations of automatic gap-filling.
2. In the current version of Kbase, there are three choices for an Annotation application: RASTtk, Prokka and DRAM, which may result in different functional annotations. When building a GEM from a genome sequence, it is recommended to try all three algorithms and compare them in terms of number of genes with unknown functions, and other quality indicators. In the case reported here, we found that the RASTtk app gave us the best result in terms of completeness of the metabolic network when compared to other GEMs, such as that of *E. coli*.
3. The gap-filling step is dependent on the growth medium chosen, because, in adding reactions to the GEM, it ensures that the model produces biomass given the current ensemble of environmentally available nutrients. Therefore, on rich media (such as LB), gap filling algorithms will likely not need to add reactions that produce essential metabolites because they are supplied in the medium. In our case study, we used a minimal medium (with glucose as a carbon source), leading to a model with a relatively large number of added reactions.
4. The final model generated by Kbase can be downloaded in different formats. Since COMETS uses the Cobra systems biology toolbox to handle metabolic models, it supports different formats. In this case, we used the SBML format, but a JSON file would have also worked well.

### Notes on COMETS simulations of a *P. simiae* colony propagation with and without chemotaxis to alanine

1. In order to take into account non-growth associated maintenance costs, metabolic models are often edited to incorporate a maintenance reaction that hydrolyzes ATP without coupling to any other reaction. The flux through this reaction is generally set to a fixed value, which needs to be determined experimentally, or inferred from literature data. For the purpose of the current analysis we set the maintenance flux to zero (see Methods 2.4).
2. Note that as described in step 2.7., here we use only the linear diffusion model. This is implemented by setting (in Eq. 1) *D*_0_ = 10^−7^*cm*^2^*/s, D*_*k*_ = 0, *k* = 0, *n* = 1, and *K* = 0. The diffusion model can be modified to add nonlinearities by modifying these parameters and can be modulated by a Hill function to prevent diffusion of non-growing biomass (see [15] for more details). A similar Hill function can be used for modulating the chemotaxis term as well.
3. Note that different GEMs may use different naming conventions for compounds. Some of the two most common ones are those defined by Kbase/Model Seed [43] and the BiGG database [44]. COMETS can handle both naming conventions. If one uses a model with the Model Seed naming convention (“cpd”), then all metabolites must begin with “cpd” followed by the compound ID. Compounds named according to the BiGG database have IDs that reflect the metabolite name (e.g. “h2o”). For successful simulations it is crucial to match the metabolite ID with the name from the GEM when setting boundary conditions.
4. The metabolite amounts per spatial location (pixel) of the simulation grid are set in Methods 2.11. It is important to remember that for each metabolite this boundary condition is defined as a net amount (with units of mmol per grid pixel) rather than a concentration. This is done to avoid confusion, given that the grid is 2D, and therefore there is no volume unambiguously associated with each pixel. If a metabolite is considered unlimiting (e.g. O2 under aerobic condition), its abundance is set to a very high number, typically 1000 mmol.
5. The chemoattractant-specific parameters for the chemotaxis model are set in 2.11 with the function `layout.set_chemotaxis(0, “cpd00035_e0”, 0.0001, 1e-6)`. Here the first parameter is the ID number of the model. In this case we have only one model, so it is set to zero. The second parameter is the name of the chemoattractant in the model. In this case, the ID of alanine in the model is `cpd00035_e0`. The third parameter is the chemotaxis coefficient *χ*, and the last one is the sensitivity *a*_*m*_ as specified in equations (1) and (2). To switch from chemoattractant to chemorepellent, one needs to change the sign of *χ*.
6. The diffusivity of the metabolites can be either globally set the same for all metabolites, or for each metabolite separately. More info can be found in Dukovski et al. (2021) [14].
7. The parameters of the simulation are organized in a class on their own. The time step and linear grid pixel size need to be chosen in a way that produces a numerically stabile simulation. A time step that is too large, in combination with a grid pixel size that is too small will result in numerical instability. Guidelines for appropriate choices can be found in Dukovski et al. (2021) [14].
8. The maximum uptake rate for a metabolite import flux in COMETS is generally computed as a function of the metabolite concentration using the Michaelis-Menten formula *V* = *V*_*max*_[*c*]*/*([*c*] + *K*_0_), where [*c*] is the concentration of the metabolite, and *V*_*max*_ and *K*_0_ are parameters of the model. These parameters can be either globally set equal for all metabolites, or separately, specifically for each metabolite. More information can be found in Dukovski et al. (2021) [14].
9. The main outputs of a COMETS simulation are total integrated biomass (over the whole space), as well as local biomass, extracellular metabolite concentrations and genome-scale fluxes for every location, time step and organism. They are saved as Pandas data frames in the simulation class. They can be accessed as `simulation.total_biomass, simulation.biomass, simulation.media` and `simulation.fluxes_by_species`. If the parameters to save them into files are set to True, the corresponding results will be printed out into files that can be analyzed independent of the simulation script.

### Notes for Visualization and analysis of the simulation output

1. The scripts in this section are specific to this simulation layout. They can easily be customized for different layouts and simulation parameters.
2. It is important to note that the file reading functions are specific to the format of the output files. The output of a COMETS simulation can be saved by the COMETS utilities, if the save output parameters are set to True. However, one can save the output by using the Pandas utility, `simulation.biomass.to_csv(‘biomass.csv’)` for example. In that case, instead of using the load functions presented here, one can load the file directly in Pandas with its utility `pandas.read_csv` and perform custom analysis.

## Acknowledgements

We would like to thank Dr. Kirill Korolev for his advice on modelling chemotaxis, Dr. Konrad Herbst for his help in agar experiments, and Alvin Lu and Suraj Prabhu for their help with the genome-scale model. This work was partially supported by the U.S. Department of Energy, Office of Science, Office of Biological & Environmental Research through the Microbial Community Analysis and Functional Evaluation in Soils SFA Program (m-CAFEs) under contract number DE-AC02-05CH11231 to Lawrence Berkeley National Laboratory; the National Science Foundation (NSF-BSF 2246707 and the NSF Center for Chemical Currencies of a Microbial Planet, Award Number 2019589); the Human Frontiers Science Program (RGP0060/2021); and the Boston University Biological Design Center Kilachand Multicellular Design Program.

## References

[1] M. A. Matilla and T. Krell, “The effect of bacterial chemotaxis on host infection and pathogenicity,” FEMS Microbiol Rev, vol. 42, no. 1, 2018, doi: 10.1093/femsre/fux052.

[2] P. Kundu, E. Blacher, E. Elinav, and S. Pettersson, “Our Gut Microbiome: The Evolving Inner Self,” Cell, vol. 171, no. 7, pp. 1481–1493, Dec. 2017.

[3] B. E. Scharf, M. F. Hynes, and G. M. Alexandre, “Chemotaxis signaling systems in model beneficial plant-bacteria associations,” Plant Mol. Biol., vol. 90, no. 6, pp. 549–559, Apr. 2016.

[4] J. M. Keegstra, F. Carrara, and R. Stocker, “The ecological roles of bacterial chemotaxis,” Nat. Rev. Microbiol., vol. 20, no. 8, pp. 491–504, Aug. 2022.

[5] W. Shou, S. Ram, and J. M. G. Vilar, “Synthetic cooperation in engineered yeast populations,” Proc. Natl. Acad. Sci. U. S. A., vol. 104, no. 6, pp. 1877–1882, Feb. 2007.

[6] J. A. Vorholt, C. Vogel, C. I. Carlström, and D. B. Müller, “Establishing Causality: Opportunities of Synthetic Communities for Plant Microbiome Research,” Cell Host Microbe, vol. 22, no. 2, pp. 142–155, 2017, doi: 10.1016/j.chom.2017.07.004.

[7] J. Kehe et al., “Massively parallel screening of synthetic microbial communities,” Proc. Natl. Acad. Sci. U. S. A., vol. 116, no. 26, pp. 12804–12809, Jun. 2019.

[8] O. S. Venturelli et al., “Deciphering microbial interactions in synthetic human gut microbiome communities,” Mol. Syst. Biol., vol. 14, no. 6, p. e8157, Jun. 2018.

[9] N. I. Johns, T. Blazejewski, A. L. Gomes, and H. H. Wang, “Principles for designing synthetic microbial communities,” Curr. Opin. Microbiol., vol. 31, pp. 146–153, Jun. 2016.

[10] T. Grosskopf and O. S. Soyer, “Synthetic microbial communities,” Curr. Opin. Microbiol., vol. 18, no. 100, pp. 72–77, Apr. 2014.

[11] F. D. Andreote and M. de C. Pereira E Silva, “Microbial communities associated with plants: learning from nature to apply it in agriculture,” Curr. Opin. Microbiol., vol. 37, pp. 29–34, Jun. 2017.

[12] J. M. Morrison, K. D. Baker, R. M. Zamor, S. Nikolai, M. S. Elshahed, and N. H. Youssef, “Spatiotemporal analysis of microbial community dynamics during seasonal stratification events in a freshwater lake (Grand Lake, OK, USA),” PLoS One, vol. 12, no. 5, p. e0177488, May 2017.

[13] H. Feng et al., “Chemotaxis of Beneficial Rhizobacteria to Root Exudates: The First Step towards Root-Microbe Rhizosphere Interactions,” Int. J. Mol. Sci., vol. 22, no. 13, Jun. 2021.

[14] I. Dukovski and others, “A metabolic modeling platform for the computation of microbial ecosystems in time and space (COMETS),” Nat. Protoc., vol. 16, pp. 5030–5082, 2021.

[15] I. Dukovski, L. Golden, J. Zhang, M. Osborne, D. Segrè, and K. S. Korolev, “Biophysical metabolic modeling of complex bacterial colony morphology,” Cell Systems, vol. 16, Issue 8, 101352, Aug. 20, 2025.

[16] M. W. Covert et al., “Metabolic modeling of microbial strains in silico,” Trends Biochem. Sci., vol. 26, no. 3, pp. 179–186, Mar. 2001.

[17] J. S. Edwards, M. Covert, and B. Palsson, “Metabolic modelling of microbes: the flux-balance approach,” Environ. Microbiol., vol. 4, no. 3, pp. 133–140, 2002.

[18] B. Ø. Palsson, Systems Biology: Properties of Reconstructed Networks. Cambridge University Press, 2006.

[19] N. D. Price, J. L. Reed, and B. Ø. Palsson, “Genome-scale models of microbial cells: evaluating the consequences of constraints,” Nat. Rev. Microbiol., vol. 2, no. 11, pp. 886–897, Nov. 2004.

[20] J. D. Orth, I. Thiele, and B. Ø. Palsson, “What is flux balance analysis?,” Nat Biotechnol, vol. 28, pp. 245–248, 2010.

[21] B. Ø. Palsson, “Systems Biology: Properties of Reconstructed Networks,” Cambridge University Press, 2006.

[22] M. A. Henson and T. J. Hanly, “Dynamic flux balance analysis for synthetic microbial communities,” IET Syst. Biol., vol. 8, no. 5, pp. 214–229, Oct. 2014.

[23] J. Chen, J. A. Gomez, K. Höffner, P. I. Phalak Poonam and Barton, and M. A. Henson, “Spatiotemporal modeling of microbial metabolism,” BMC Syst. Biol., vol. 10, p. 21, Mar. 2016.

[24] R. Mahadevan, J. S. Edwards, and F. J. Doyle 3rd, “Dynamic flux balance analysis of diauxic growth in Escherichia coli,” Biophys. J., vol. 83, no. 3, pp. 1331–1340, Sep. 2002.

[25] K. Höffner, S. M. Harwood, and P. I. Barton, “A reliable simulator for dynamic flux balance analysis,” Biotechnol. Bioeng., vol. 110, no. 3, pp. 792–802, Mar. 2013.

[26] W. R. Harcombe and others, “Metabolic resource allocation in individual microbes determines ecosystem interactions and spatial dynamics,” Cell Rep., vol. 7, pp. 1104–1115, 2014.

[27] N. Wadhwa and H. C. Berg, “Bacterial motility: machinery and mechanisms,” Nat. Rev. Microbiol., vol. 20, no. 3, pp. 161–173, Mar. 2022.

[28] M. Miyata et al., “Tree of motility - A proposed history of motility systems in the tree of life,” Genes Cells, vol. 25, no. 1, pp. 6–21, Jan. 2020.

[29] R. Karmakar, “State of the art of bacterial chemotaxis,” J Basic Microbiol, vol. 61, no. 5, pp. 366–379, May 2021, doi: 10.1002/jobm.202000661.

[30] D. Lauffenburger, R. Aris, and K. H. Keller, “Effects of random motility on growth of bacterial populations,” Microb. Ecol., vol. 7, no. 3, pp. 207–227, Sep. 1981.

[31] I. Sampedro, R. E. Parales, T. Krell, and J. E. Hill, “Pseudomonas chemotaxis,” FEMS Microbiol. Rev., vol. 39, no. 1, pp. 17–46, Jan. 2015.

[32] M. J. Tindall, P. K. Maini, S. L. Porter, and J. P. Armitage, “Overview of mathematical approaches used to model bacterial chemotaxis II: bacterial populations,” Bull. Math. Biol., vol. 70, no. 6, pp. 1570–1607, Aug. 2008.

[33] E. F. Keller and L. A. Segel, “Initiation of slime mold aggregation viewed as an instability,” J Theor Biol, vol. 26, no. 3, pp. 399–415, 1970, doi: 10.1016/0022-5193(70)90092-5.

[34] E. F. Keller and L. A. Segel, “Model for chemotaxis,” J. Theor. Biol., vol. 30, no. 2, pp. 225–234, Feb. 1971.

[35] E. F. Keller and L. A. Segel, “Traveling bands of chemotactic bacteria: a theoretical analysis,” J Theor Biol, vol. 30, no. 2, pp. 235–248, 1971, doi: 10.1016/0022-5193(71)90051-8.

[36] M. Burger, M. Di Francesco, and Y. Dolak-Struss, “The Keller–Segel model for chemotaxis with prevention of overcrowding: Linear vs. nonlinear diffusion,” SIAM J. Math. Anal., 2006.

[37] M. P. Brenner, L. S. Levitov, and E. O. Budrene, “Physical Mechanisms for Chemotactic Pattern Formation by Bacteria,” Biophys., vol. 74, no. 4, pp. 1677–1693, Apr. 1998, doi: 10.1016/S0006-3495(98)77880-4.

[38] T. Bhattacharjee, D. B. Amchin, R. Alert, J. A. Ott, and S. S. Datta, “Chemotactic smoothing of collective migration,” Elife, vol. 11, Mar. 2022, doi: 10.7554/eLife.71226.

[39] T. Curk, D. Marenduzzo, and J. Dobnikar, “Chemotactic Sensing towards Ambient and Secreted Attractant Drives Collective Behaviour of E. coli,” PLoS One, vol. 8, no. 10, p. e74878, Oct. 2013, doi: 10.1371/journal.pone.0074878.

[40] J. Cremer, T. Honda, Y. Tang, J. Wong-Ng, M. Vergassola, and T. Hwa, “Chemotaxis as a navigation strategy to boost range expansion,” Nature, vol. 575, no. 7784, pp. 658–663, Nov. 2019, doi: 10.1038/s41586-019-1733-y.

[41] A. V Narla, J. Cremer, and T. Hwa, “A traveling-wave solution for bacterial chemotaxis with growth,” Proceedings of the National Academy of Sciences, vol. 118, no. 48, p. e2105138118, 2021, doi: 10.1073/pnas.2105138118.

[42] C. M. J. Pieterse et al., “Pseudomonas simiae WCS417: star track of a model beneficial rhizobacterium,” Plant Soil, vol. 461, no. 1–2, pp. 245–263, Apr. 2021, doi: 10.1007/s11104-020-04786-9.

[43] A. P. Arkin et al., “KBase: The United States department of energy systems biology knowledgebase,” Jul. 06, 2018, Nature Publishing Group. doi: 10.1038/nbt.4163.

[44] Z. A. King et al., “BiGG Models: A platform for integrating, standardizing and sharing genome-scale models,” Nucleic Acids Res, vol. 44, no. D1, pp. D515–D522, 2016, doi: 10.1093/nar/gkv1049.

